# Phosphorylation controls RNA binding and transcription by the influenza virus polymerase

**DOI:** 10.1101/2020.02.10.942318

**Authors:** Anthony R. Dawson, Gary M. Wilson, Elyse C. Freiberger, Arindam Mondal, Joshua J. Coon, Andrew Mehle

## Abstract

The influenza virus polymerase transcribes and replicates the viral genome. The proper timing and balance of polymerase activity is important for successful replication. We previously showed that phosphorylation regulates genome replication by controlling assembly of the replication machinery (Mondal, et al. 2017). However, it remained unclear whether phosphorylation directly regulated polymerase activity. Here we identified polymerase phosphosites that control its function. Mutating phosphosites in the catalytic subunit PB1 altered polymerase activity and virus replication. Biochemical analyses revealed phosphorylation events that disrupted global polymerase function by blocking the NTP entry channel or preventing RNA binding. We also identified a regulatory site that split polymerase function by specifically suppressing transcription. These experiments show that host kinases phospho-regulate viral RNA synthesis directly by modulating polymerase activity and indirectly by controlling assembly of replication machinery. Further, they suggest polymerase phosphorylation may bias replication versus transcription at discrete times or locations during the infectious cycle.

## Introduction

All RNA viruses encode machinery both to express viral transcripts and to replicate genomes. Negative sense RNA viruses must first transcribe using virally-encoded RNA-dependent RNA polymerases (RdRPs) that are packaged into virions. The viral RdRP subsequently replicates the genome, often with the help of protein products from the recently produced mRNA. Regulating the balance and timing of transcription and replication is crucial for successful infection.

Viruses employ diverse strategies to control the abundance of virally-derived RNAs. Many RNA viruses rely on RdRP co-factors whose activity is dynamically regulated by post-translational modifications. For example, the Ebola virus polymerase is regulated by the viral transcription factor VP30. VP30 promotes transcription, whereas phosphorylation of VP30 results in its exclusions from transcription complexes favoring genome replication (Biedenkopf et al., 2016). A similar strategy is employed by Marburg virus (Tigabu et al., 2018). Dynamic phosphorylation of the M2-1 protein from respiratory syncytial virus regulates viral transcription. M2-1 is a transcriptional processivity factor whose function is proposed to require cycles of phosphorylation and dephosphorylation by cellular enzymes (Cartee and Wertz, 2001; Richard et al., 2018). Phosphorylation also regulate global RNA synthesis. The phosphoprotein (P) from vesicular stomatitis virus is a polymerase co-factor. Phosphorylation on the N-terminus of P is important for transcription and replication (Chen et al., 2013; Mondal et al., 2014). The dynamic and fully reversible nature of phosphorylation enables localized and temporal control of viral proteins and may help progression through the infectious cycle. Phosphorylation of polymerase co-factors is thus a common strategy to regulate transcription and replication. However, influenza A virus and other members of *Orthomyxoviridae* do not encode polymerase co-factors and it remains unclear how their polymerases are regulated.

Influenza A virus contains eight negative-sense RNA genome segments packaged into virions as ribonucleoprotein (RNP) complexes. RNPs are double helical flexible rod-like structures containing the viral genome coated by oligomeric nucleoprotein (NP) and bound at both ends by the viral polymerase (Arranz et al., 2012; Klumpp et al., 1997; Moeller et al., 2012; Pons et al., 1969). The viral polymerase is a heterotrimeric complex composed of the PB1, PB2 and PA subunits. Immediately following uncoating, RNPs are trafficked to the nucleus where synthesis of all virally-derived RNA occurs (Herz et al., 1981; Jackson et al., 1982). Infection initiates with a pioneering round of transcription from the incoming RNPs. The viral polymerase performs cap-snatching where a short capped oligonucleotide derived from the host is used to prime transcription (Bouloy et al., 1978; Plotch et al., 1979). Unlike transcription, replication initiates in a primer-independent fashion to create a positive-sense intermediate (cRNA) that serves as a template for vRNA production (Hay et al., 1977). Replication requires concomitant assembly into newly formed RNPs to stabilize the viral genome (Vreede et al., 2004). Newly formed vRNPs can either be packaged into virions or serve as templates for additional rounds of transcription or replication.

The processes regulating polymerase activity are not fully defined. Some regulation simply requires the production of specific viral proteins or RNAs. The stable products from incoming RNPs are viral mRNAs, even though incoming RNPs are capable of making cRNA as well. Replication occurs later when newly synthesized NP is able to coat cRNA genomes and protect them from degradation (Vreede et al., 2004). Newly synthesized viral polymerase binds nascent RNA products and interacts with cRNP-bound polymerases to stimulate production of full-length vRNA (Fodor and te Velthuis, 2019). The nuclear export protein (NEP) and small viral RNAs (svRNA), both of which are made at later stages of infection, further bias the polymerase to replication (Perez et al., 2010; Robb et al., 2009) Infection also induces broad changes in signaling cascades, and multiple host and viral proteins are regulated by post-translational modifications during influenza virus infection (Dawson and Mehle, 2018). Like many other RNA viruses, phosphorylation of viral proteins plays a key role in regulating the influenza virus replication machinery. We have previously shown that phosphorylation of NP regulates *de novo* RNP assembly (Mondal et al., 2015, 2017). The protein kinase C (PKC) family, and PKCδ in particular, phosphorylates NP at its homotypic interface to block NP oligomerization. This is proposed to create a pool of monomeric NP that is subsequently licensed for oligomerization by a cellular phosphatase, possibly CDC25B (Cui et al., 2018).

PKCδ phospho-regulates NP oligomerization and by extension the ability of the polymerase to replicate the viral genome (Mondal et al., 2017). These studies also provided intriguing data suggesting that the polymerase may also be phosphorylated. Whether phosphorylation directly regulates polymerase activity is unclear. Here we extensively map phosphorylation sites on the polymerase subunit PB1 and characterize their function. PB1 is the structural and catalytic core of the enzyme, and we define PB1 phospho-sites that inhibit RNA synthesis by blocking global catalytic function or genomic RNA binding. We also identified a regulatory site that split the function of the polymerase; mimicking phosphorylation at PB1 S673 suppressed transcription without altering genome replication. Viruses encoding phospho-ablative mutants at these positions displayed altered replication kinetics, whereas phospho-mimetic mutants did not replicate. These data demonstrate that phosphorylation directly regulates viral polymerase activity and may provide a mechanism to bias populations of polymerase towards replication or transcription.

## Results

### Phosphorylation alters activity of influenza virus polymerase

We had previously shown that PKCδ regulates RNP assembly by modifying NP and preventing premature NP oligomerization (Mondal et al., 2017). This work also revealed slower migrating species of the PB2 polymerase subunit that raised the possibility that the polymerase itself was phosphorylated in the presence of PKCδ. To test this possibility, we assessed the migrations patterns of PB2 before or after phosphatase treatment. We sought to study direct effects on the viral polymerase, but this cannot be done in the context of an RNP as PKCδ regulates NP function. We eliminated this confounder by using a short vRNA template (vNP77) that does not require NP for replication or transcription, and thus decouples RNP assembly from RNA synthesis activities (Turrell et al., 2013). The viral polymerase was expressed in cells with constitutively active PKCδ and vNP77 and immuno-purified samples were analyzed by western blot (Figure 1A). Slower migrating species were detected for PB2, confirming our prior results. Treating samples with phosphatase collapsed these species into a single band migrating at the expected position for PB2, suggesting that PB2 is phosphorylated. A shorter exposure of the same gel confirmed equivalent loading of PB2.

**Figure 1:**
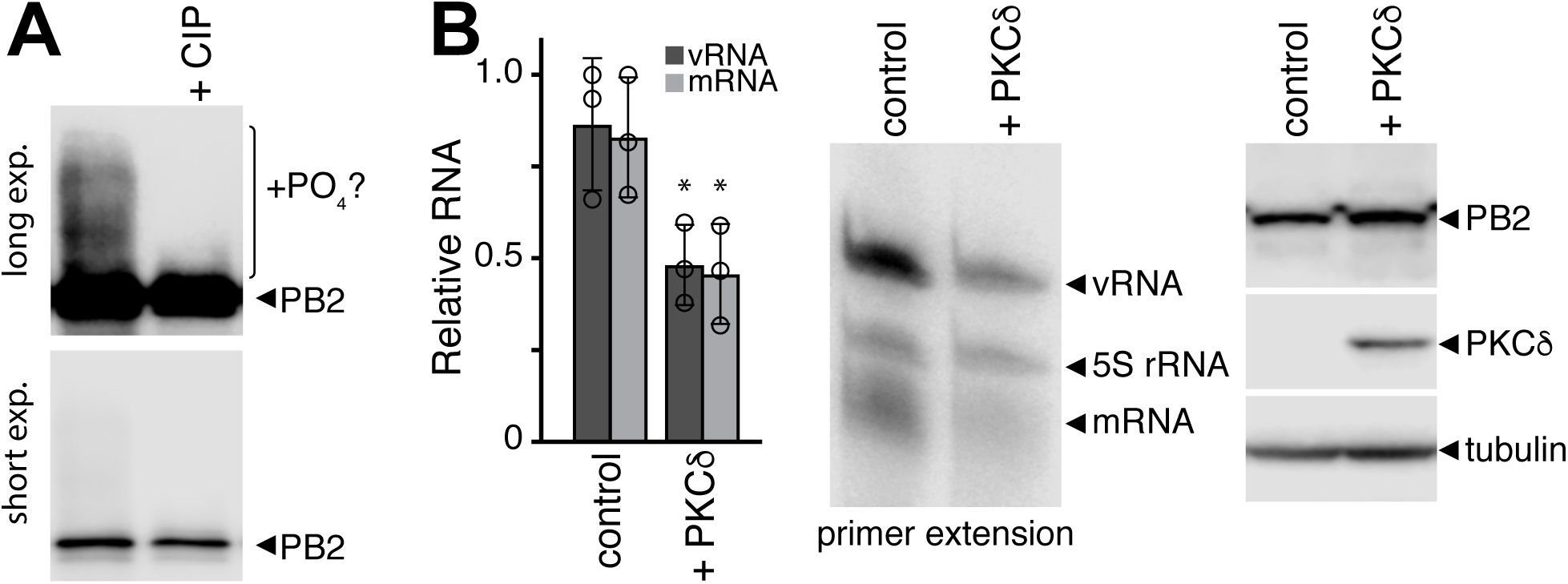
PKCδ stimulates polymerase phosphorylation and inhibits polymerase activity. **A.** The viral polymerase proteins and the mini-gene vNP77 were expressed in cells with a constitutively active form of PKCδ. The polymerase was immunopurified, mock treated or incubated with calf-intestinal phosphatase (CIP), and analyzed by western blot. Long and short exposures of the same blot are shown. A CIP-sensitive species that may indicate phosphorylated PB2 is indicated. **B.** Primer extension assays were performed on RNA extracted from cells expressing the viral polymerase, vNP77 and constitutively active PKCδ or an empty vector control. Viral replication (vRNA) and transcription (mRNA) were quantified, normalized to the 5S rRNA internal control, and presented relative to polymerase without PKCδ (mean of n=3 ± sd; * = Student’s t-test P < 0.05). Representative primer extension data are shown with western blots to confirm protein expression.

We then asked whether PKCδ expression affects RNA synthesis activities of the polymerase. The viral polymerase and vNP77 were expressed in cells in the presence or absence of constitutively-active PKCδ. Primer extension analysis of RNA extracted from these samples quantified transcription (mRNA) and replication (vRNA) products. Co-expression of PKCδ significantly impaired production of viral transcripts and replication products, without altering viral protein levels (Figure 1B). Together, these data indicate that the viral polymerase is phosphorylated and these modifications may alter intrinsic polymerase activity.

### The polymerase core is phosphorylated at highly conserved sites

Given its potential to regulate polymerase activity, we performed a series of complementary experiments to extensively characterize polymerase phosphorylation (Figure 2A). We repeated experiments where phosphorylation was shown at affect polymerase function by expressing the viral polymerase, vNP77 and activated PKCδ in 293T cells. Polymerase was immuno-purified and analyzed phospho-peptide mass spectrometry (MS) in two independent experiments. In parallel, we analyzed samples from infected cells. Polymerase phosphorylation status may vary across multiple rounds of infection; therefore, we collected samples from low and high MOI infections performed in A549 cells and analyzed whole-cell lysate. We also allowed for the possibility that phosphorylation patterns change throughout a single infection by analyzing RNP immunoprecipitations from both individual and pooled time points from synchronized infections. The amount of each sample used in the pooled lysate was adjusted to approximate similar levels of viral protein for all time points. These approaches allowed for high-confidence identification of phosphorylation sites on the viral polymerase (Supplemental Table 1).

**Figure 2:**
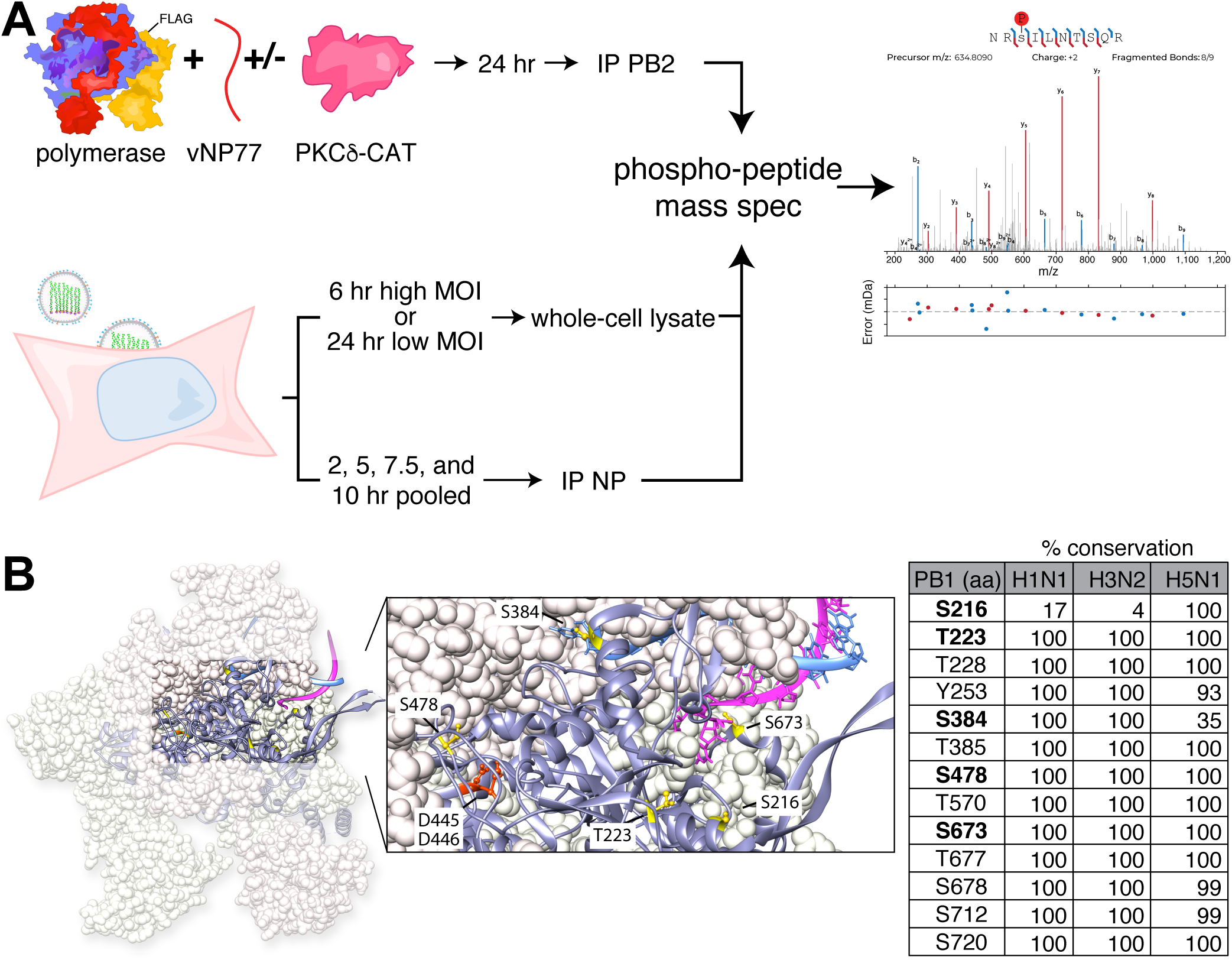
The catalytic core of the viral polymerase is phosphorylated during infection. **A.** Experimental design to detect phosphorylation of an active polymerase in transfected 293T cells or infected A549 cells. Samples were prepared as whole-cell lysate or immuno-purified proteins prior to phospho-peptide mass spectrometry. A representative spectra is show. See Supplemental Table 1 for all identified sites. **B.** Phospho-sites on PB1 surround the catalytic core and template entry. The location of PB1 phospho-sites characterized in this study are modeled in yellow on PDB 4WSB (Reich et al., 2014). The motif C residues D445/D446 in the catalytic site are in orange, 5’ vRNA is blue, and 3’ vRNA is magenta. The space-filled representation of PA and PB2 are shown as light pink and light yellow, respectively. Most phospho-sites are conserved among circulating human influenza virus strains and highly pathogenic H5N1 viruses. Percentage of sequences where the indicated residue is a serine, threonine, or tyrosine are shown.

We focused our analysis on PB1, the subunit that catalyzes RNA synthesis. A total of 13 phosphorylated residues were identified on PB1 in all experimental conditions, of which 8 were also detected in the context of infection (Supplemental Table 1). Phosphorylation occurred primarily on threonine and serine residues, with only one phospho-tyrosine identified. Most of the phospho-sites are highly conserved in human H1N1 and H3N2 strains and highly pathogenic H5N1 strains (Figure 2B). Some of the identified phosphorylation sites overlapped between our infected and transfected cells (S216, T223, S673), whereas others were identified only during infection (T228, Y253, T570, S712, S720) or when polymerase was co-expressed with PKCδ (S478, S384). We placed high priority on the four sites that we identified in at least two separate experiments: PB1 S216, T223, S384, and S673. These phospho-sites could be broadly categorized into those that are proximal to the catalytic center (S216) or the template entry channel (T223, S384 and S673) (Figure 2B). We also included PB1 S478 given its close proximity to the catalytic center. Phosphorylation at PB1 T223 was identified during infection, confirming and extending the importance of prior work that had identified this phospho-site from transfected cells (Hutchinson et al., 2012). None of the other phospho-sites have been previously reported.

### Phosphorylation status of PB1 impacts the ability of the polymerase to produce RNA and infectious virions

To assess the biological relevance of these phosphorylation events, we attempted to rescue influenza virus encoding PB1 mutants harboring phospho-mimetic aspartic acid (D) or phospho-ablative alanine (A) mutations. All tested phospho-ablative PB1 constructs produced virus, whereas phospho-mimetic mutants at PB1 T223, S478 and S673 failed to yield virus despite multiple attempts. For PB1 mutants that support production of infectious virus, we measured viral gene expression during single-cycle infection of A549 cells (Figure 3A). Phospho-ablative mutants produced similar amounts of NP mRNA compared to WT, with the exception of PB1 T223A that had decreased transcription and PB1 S673A that exhibited a significant 5-fold increase in gene expression. Phospho-mimetic mutants displayed more subtle phenotypes, with PB1 S216D slightly above and PB1 S384D was slightly below WT levels. These viruses were then assayed in a multi-cycle replication assay (Figure 3B). All viruses exhibited roughly similar replication kinetics at 12 and 24 hpi. However, PB1 T223A plateaued at peak titers ∼10-fold lower than WT and the titers of PB1 S478A and S673A rapidly declined at 72 and 96 hpi to yield final titers ∼10-fold lower than WT. PB1 harboring phospho-mimetics at position S216 and S384 yielded virus that replicates similar to WT virus despite producing disparate levels of NP transcripts (Figure 3A-B). Constitutive phosphorylation at PB1 T223, S478, and S673 is incompatible with production of infectious virus, whereas the complete loss of phosphorylation at positions PB1 T223 and S673 also disrupts transcription and viral replication. These data suggest that differential phosphorylation of PB1 is important for successful infection.

**Figure 3:**
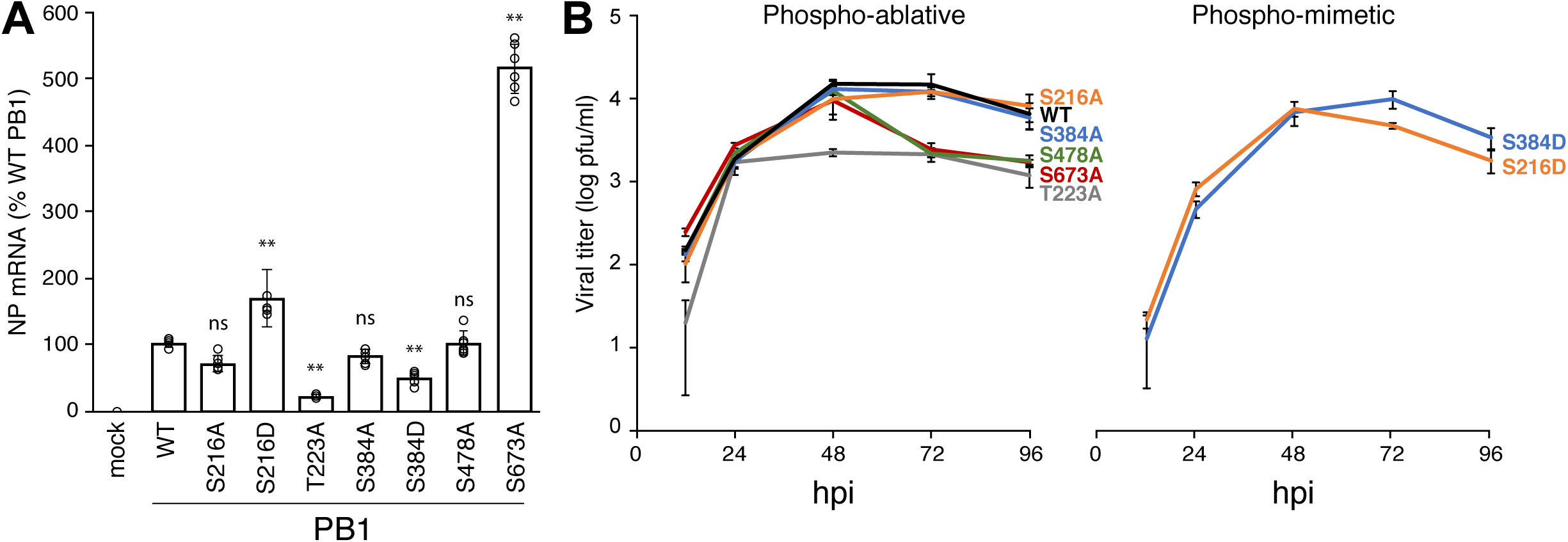
PB1-mutant viruses identify phosphorylation sites that impact polymerase activity and viral replication. **A.** PB1 phosphorylation both inhibits and enhances viral transcription. Single-cycle infections with PB1 phospho-mutant viruses were performed in A549 cells (MOI of 0.5 for 8h). RNA from infected cells was subject to qRT-PCR to detect NP and GAPDH mRNA. Fold changes (ΔΔCT) were determined in triplicate from 2 independent infections. (± SEM; ** = P <0.01 for one-way ANOVA with Dunnett’s post hoc compared to WT). **B.** Multicycle replication kinetics of phospho-mutant viruses. A549 cells were infected at an MOI of 0.001. Viral titers were measured 12, 24, 48, 72, 96 hpi via plaque assay on MDCK cells. (mean of n=3 ± sd). P < 0.01 for one-way ANOVA at each time point. Statistics for ANOVA with Dunnett’s post hoc pair-wise comparisons to WT are in Supplemental Table 2.

Replication assays revealed PB1 phospho-residues important for the infectious cycle. We next focused solely on polymerase activity by performing primer extension assay for polymerases containing WT or phospho-mutant PB1 (Figure 4A, Figure 4 Figure Supplement 1). WT polymerase produced significant amounts of viral mRNA, the replication intermediate cRNA, and vRNA indicating successful transcription and replication of the input genome. Phospho-mimetic mutants that failed to produce infectious virus also displayed defects in RNA synthesis. Polymerases with PB1 S223D or S478D exhibit profound defects with only background input vRNA levels and no detectable transcription or replication products. These results mirrored those obtained with a catalytically dead PB1 D445A/D446A mutant (PB1a) (Vreede et al., 2004). Remarkably, PB1 S673D replicated viral RNA, but showed a severe reduction in mRNA. Transcriptional defects for PB1 S673D were as strong as the previously described transcriptional mutant PB1 K669A/R670A (Figure 4A, Figure 4 Figure Supplement 1)(Kerry et al., 2008). These data suggest that phosphorylation at PB1 S673 biases plus-sense RNA synthesis away from transcription and towards replication. PB1 S216D, S384D, and S673A generated transcription and replication products similar to WT PB1. Some of the differences in transcription detected during single-cycle infection (Figure 3A) were not fully recapitulated in primer extension assays (Figure 4A). This could be explained by the simplified nature of primer extension assays that lack viral factors that may modulate RNA production during infection (Robb et al., 2009).

**Figure 4:**
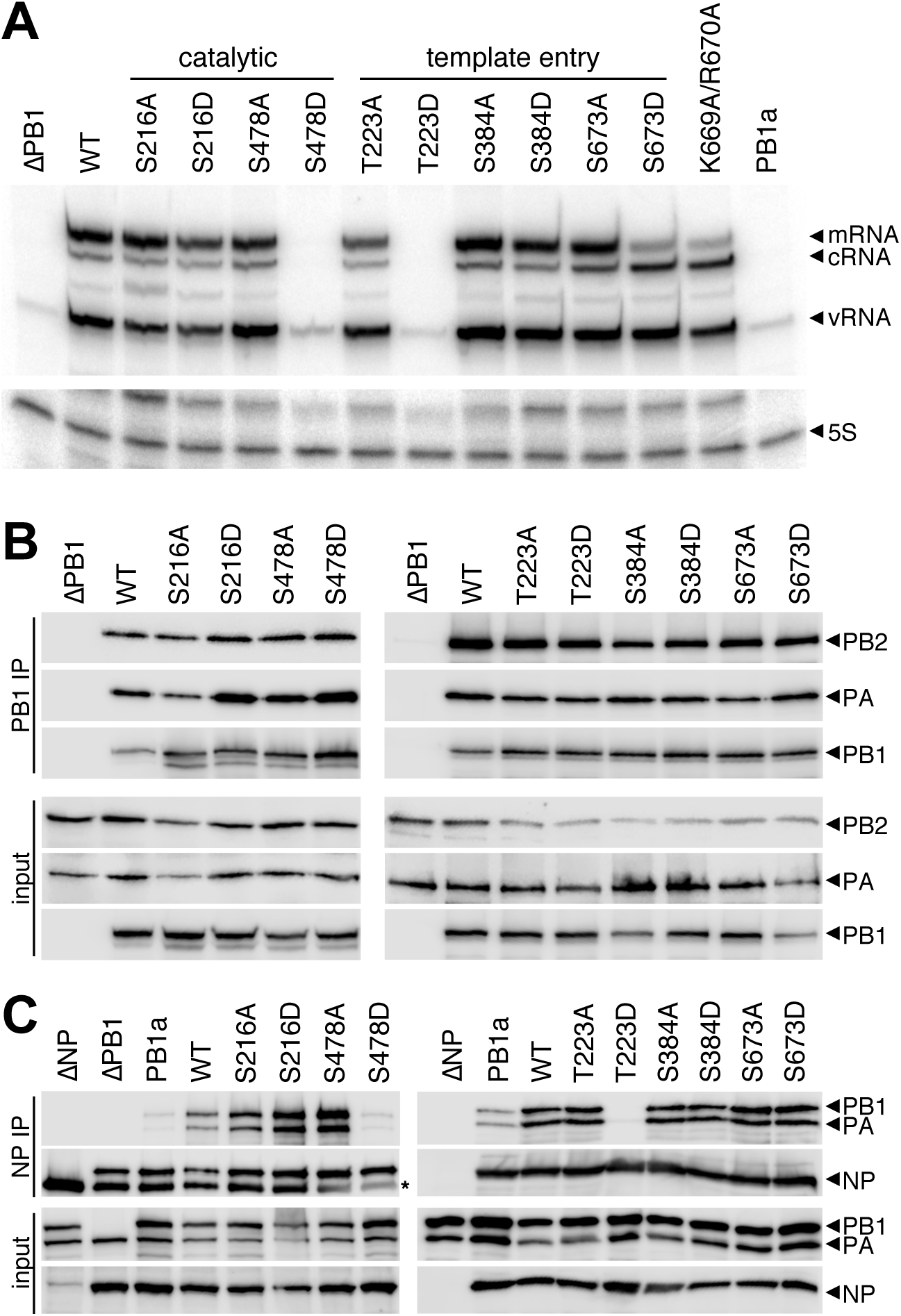
PB1 phospho-mutants are defective in RNA synthesis and RNP formation. **A.** Viral RNA synthesis was analyzed in primer extensions assays. RNA extracted from 293T cells expressing the viral polymerase, NP, and segment 6 vRNA was subject to primer extension analysis to detect transcription (mRNA) and replication (cRNA, vRNA) products. Primer extension of 5S rRNA was used as an internal loading control. PB1a, catalytically-dead PB1; PB1 K669A/R670A, transcription-deficient PB1. **B.** PB1 phospho-mutants form polymerase trimers. FLAG-tagged PB1, PB2, and PA were expressed in 293T cells and cell lysates were subject to PB1-FLAG immunoprecipitation. Immunoprecipitates and input samples were probed for PB1-FLAG, PB2, and PA. C. PB1 phospho-mimetics deficient in RNA synthesis fail to generate productive RNPs. NP immunoprecipitations were performed on 293T lysates generated as in (A). Immunoprecipitates and input samples were western blotted for PB1, PA, and NP.

Polymerase activity requires multiple steps for successful replication and transcription, beginning with protein expression, trimer assembly, RNA binding, RNP assembly, and ultimately the synthesis of new RNA products (Fodor and te Velthuis, 2019). Primer extension reports on the cumulative success of this process. We therefore systematically investigated each step to identify regulatory points affected by PB1 phosphorylation. PB1 stability and polymerase trimer formation were assayed by expressing proteins in cells, immuno-purifying PB1, and probing for co-precipitating PB2 and PA (Figure 4B). All PB1 mutants expressed and formed trimers at approximately WT levels, independent of whether they were phospho-ablative or phospho-mimetic. Thus, phosphorylation of PB1 at these sites does not control trimer assembly. Additionally, as polymerase trimers form in the cell nucleus, these data imply that defects in RNA production are not due to faulty nuclear import of polymerase subunits (Deng et al., 2005). RNP assembly was next investigated by expressing RNP components in cells, immuno-precipitating NP and probing for co-precipitating polymerase (Figure 4C). Active polymerase will replicate the viral genome and amplify RNP assembly. We therefore utilized the catalytically dead PB1a to measure initial RNP formation that is independent of polymerase activity. PB1 mutants with defects in polymerase activity in primer extension assays failed to form productive RNPs, but the extent of the defect suggests different causes. PB1 T223D was completely excluded from RNPs, despite that fact that it forms trimers, suggesting phosphorylation at this position precludes incorporation into an RNP. PB1 S478D, however, assembled RNPs at low levels comparably to PB1a, indicating that RNP assembly *per se* is unaffected by this mutant. Rather, RNP assembly defects here stem from catalytic defects in the polymerase and not other steps in the process. The PB1 S673D phospho-mimetic do not alter RNP assembly, consistent with its ability to synthesize WT levels of genomic RNAs. In sum, loss of phosphorylation did not alter assembly and activity of RNPs in these assay. Conversely, mimicking constitutive phosphorylation at PB1 T223 or PB1 S478 prevented formation of productive RNPs and disrupted RNA synthesis. Finally, phosphorylation at PB1 S673 appears to toggle the viral polymerase primarily into replication mode.

### PB1 T223 phosphorylation inhibits vRNA binding and cRNA stabilization

Incoming RNPs synthesize both viral mRNA and cRNA, but cRNA is rapidly degraded; the polymerase and NP that would assemble into RNPs and protect cRNA from degradation have not yet been synthesized (Vreede et al., 2004). Whereas PB1 S478D was able to form low levels of RNPs, the complete failure of PB1 T223D was suggestive of defects in RNA binding. To test this possibility, we examined whether PB1 mutants could bind and stabilize cRNA during infection (Figure 5A). Polymerase with WT or mutant PB1 was expressed in cells prior to infection with WT virus. The oligomerization-deficient NP_E339A_ was also pre-expressed to help stabilize cRNA while focusing the assay only on RNA made from the incoming RNPs. Cells were treated with actinomycin D during infection so only pre-expressed viral proteins were present. Primer extension showed that cRNA was stabilized by WT PB1 as expected (Vreede et al., 2004). Equivalent levels of vRNA in each condition confirmed efficient infection and delivery of vRNPs in all settings (Figure 5 Supplemental Figure 1). Trimers harboring PB1 S478D, which are unable to synthesize viral RNAs (Figure 4A), still stabilized cRNA yielding levels slightly higher than WT (Figure 5A). PB1 T223A also showed a minor increase in cRNA levels. However, polymerases with PB1 T223D exhibited a significant drop in cRNA stabilization. All of the other phospho-mutant polymerases stabilized cRNA to WT levels, consistent with their ability to replicate viral RNA. This was true even for PB1 S673D, which is replication competent but produces lower levels of mRNA (Figure 4A).

**Figure 5:**
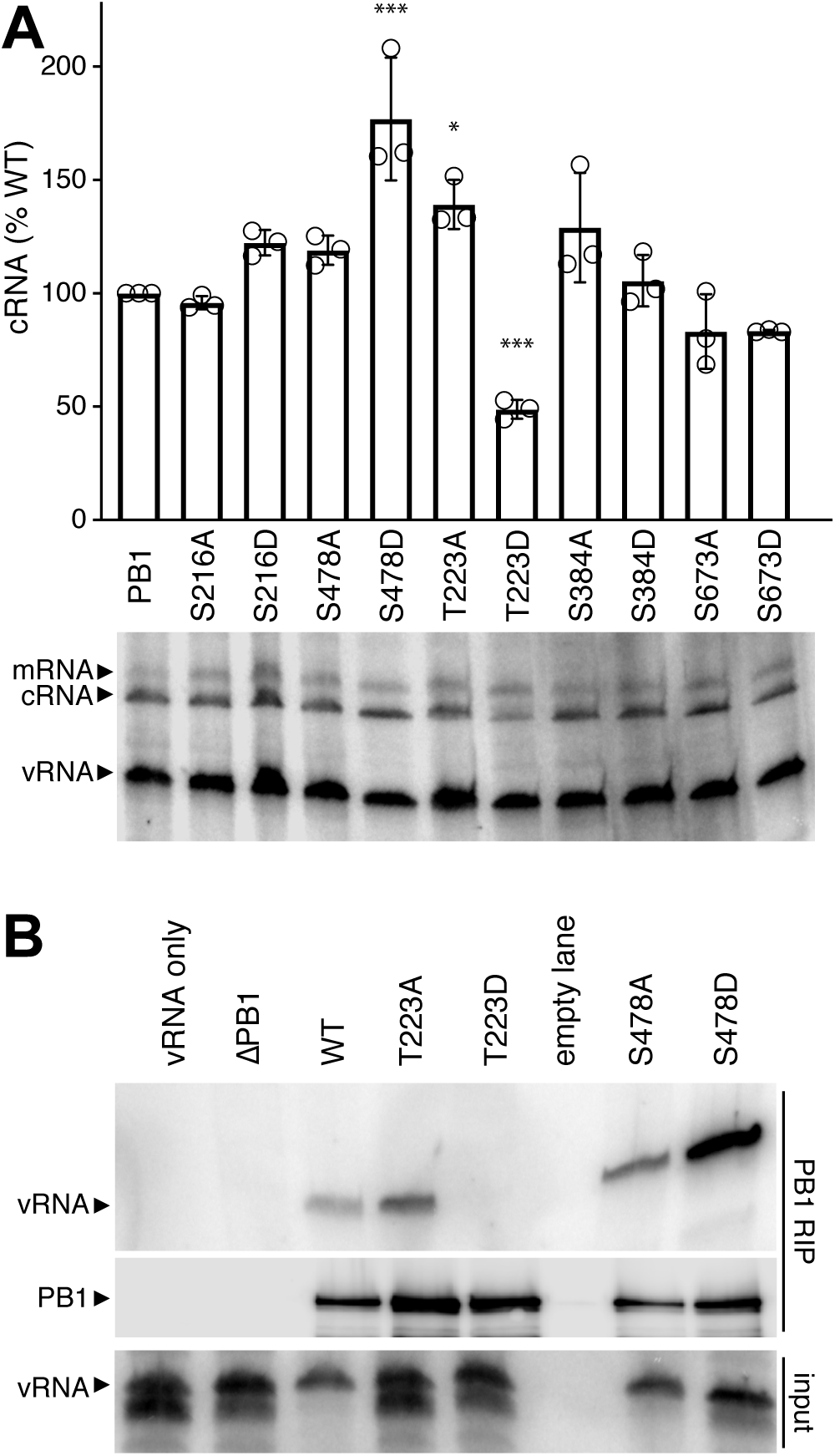
Phosphorylation at T223 inhibits cRNA stabilization and vRNA binding. **A.** PB1 phospho-mutants were tested in a cRNA stabilization assay. WT or mutant PB1 and oligomerization-deficient NP (NPE339A) were expressed in 293T cells. Cells were treated with actinomyocin D (ActD) prior to infection, RNA extraction and primer extension analysis to detect transcription (mRNA) and replication (cRNA, vRNA) products. A representative primer extension gel is shown. cRNA levels were quantified, normalized to WT, and expressed as mean ± sd. * < 0.05, ** < 0.01, *** < 0.001 = P for one-way ANOVA with Dunnett’s post hoc compared to WT. **B.** PB1 T223D fails to precipitate vRNA in an RNA-IP (RIP). PB1-FLAG, PB2, PA, and segment 6 vRNA were expressed in 293T cells. Cells were lysed and subject to FLAG immunoprecipitation. RNA extracted from immunoprecipitates and input samples was probed for the presence of segment 6 vRNA via primer extension analysis. Immunoprecipitated PB1 was confirmed via western blot.

Viral RNA promoter binding is essential for stabilizing genomic RNA, suggesting that mimicking phosphorylation at PB1 T223 interferes with RNA binding. RNA immunoprecipitation assays were performed to measure promoter binding (Figure 5B). Polymerase and segment 6 vRNA were expressed in cells and polymerase was purified by immunoprecipitating PB1. Co-precipitating vRNA was detected by primer extension. vRNA co-purified with WT PB1, but not in its absence or when all polymerase subunits were excluded from the assay. PB1 T223D completely failed to bind vRNA, even though vRNA was readily detected for PB1 T223A. PB1 S478D bound vRNA, consistent with its ability to stabilize cRNA and form low levels of RNPs. If anything, PB1 S478D showed higher binding than both WT and PB1 S478A. These data parallel those from the stabilization assays. Multiple lines of investigation identify discrete defects caused by mimicking phosphorylation. Constitutive phosphorylation at PB1 S478 appears to disrupt catalysis without affecting RNA binding or initial RNP formation. Conversely, PB1 T223D was unable to stabilize cRNA or bind vRNA, suggesting that its inability to assemble RNPs and synthesize viral RNAs arises from defects in template binding.

## Discussion

Multiple mechanisms converge to regulate the influenza polymerase and bias production of either transcripts, the replication intermediate cRNA, or genomic vRNA. Here we reveal that phosphorylation of the polymerase directly regulates its activity. The core polymerase subunit PB1 was phosphorylated at multiple conserved sites, with clusters proximal to the catalytic center or the template entry channel. Mimicking phosphorylation at PB1 S478 or T223 disrupted global polymerase function by affecting catalytic activity or template binding, respectively. By contrast, phosphorylation at PB1 S673 preferentially suppressed transcription to create a replicase form of the polymerase. In all cases, mimicking phosphorylation at these key sites blocked the production of infectious virus, whereas ablating phosphorylation led to defects in replication. These data demonstrate that phosphorylation directly controls polymerase activity by either inhibiting or regulating RNA production.

The influenza polymerase performs diverse functions as transcription peaks early in infection, followed by production of the replication intermediate cRNA, and ultimately the final replication product vRNA (Robb et al., 2009). The different functions are achieved in part by discrete conformations of the viral polymerase (Fodor and te Velthuis, 2019). One major change involves re-positioning of the 3’ end of template RNA from the surface of the polymerase to the active site (Reich et al., 2014, 2017). Two of our PB1 phospho-mutants may affect this process. Phospho-mimetics at PB1 T223 completely disrupts RNA binding and stabilization of the nascent genome, ablating polymerase activity and the production of mutant virus (Figures 4-5). Our data confirm prior identification of this phospho-site and predictions that phosphorylation at this position might alter RNA binding (Hutchinson et al., 2012; Weber et al., 2019). Whereas phosphorylation at PB1 T223 disrupts all RNA binding and activity, phosphorylation at PB1 S673 appears to differentially suppress transcription without affecting replication (Figure 4). Close inspection of viral structures reveals a potential mechanism. The 3’ end of the vRNA genome makes a pronounced turn at bases G9 and C8 as it threads through the template entry channel (Pflug et al., 2014). Flexibility at this site allows the end of the template to be directed into the active site in an initiation conformation, or to the periphery of the polymerase in a pre-initiation state (Kouba et al., 2019). Residue S673 is positioned in the crook of this turn and makes hydrogen bonds with the backbone of U7 and U10 (Reich et al., 2014). Once in the active site, the 3’ end of the template must be located in distinct positions suitable for either primer-independent replication or primer-dependent transcription (Kouba et al., 2019; Reich et al., 2017). It is possible that phosphorylation at PB1 S673 alters the trajectory of the template RNA in a way that prevents pairing with the primer or its extension, without disrupting the replication conformations or RNA binding altogether. Alternatively, basic residues K669, R670 and R672 in this region had previously been shown to be important for transcription and activating cap binding by PB2, but not replication (Kerry et al., 2008). Perhaps phosphorylation at S673 partially neutralizes their charge to reveal the same phenotype as when these basic residues were mutated to alanine. Independent of the exact molecular mechanism, phosphorylation at PB1 S673 could be a way to establish replicase-specific polymerases.

PKCδ interacts with the viral polymerase and modifies NP to control RNP assembly (Mondal et al., 2017). Here we showed that phosphorylation was identified on PB1 S478 in cells expressing active PKCδ. Mutational analysis revealed that the phospho-mimetic PB1 S478D is functionally analogous to the well-characterized PB1a allele that mutates the conserved SDD motif in the active site: PB1 S478D retains the ability to bind the viral promoter, stabilizes cRNA, and forms initial RNPs, but is catalytically inactive (Figure 4-5). S478 lies in the NTP channel of the polymerase and phosphorylation would likely interfere with NTP transit or positioning. (Kouba et al., 2019). This polymerase is catalytically inactive, but like PB1a may still impact overall polymerase output by functioning in *trans* to stimulate cRNP activity (York et al., 2013).

Our studies reveal that constitutive phosphorylation largely inhibits specific polymerase functions, whereas phospho-ablative mutants are more tolerated. Phospho-ablative mutants at PB1 T223, S478 and S673 retained polymerase function, but exhibit dysregulated abundance and altered replication profiles (Figure 3). These data suggest that balanced phosphorylation at these positions is important for normal polymerase output. The phospho-sites are all located at sites that are not immediately on the surface of the trimeric polymerase. This suggests kinases modify PB1 during its translation or assembly into the trimer. It further raises the possibility that these modifications cannot be accessed by a phosphatase and are thus static. Instead of dynamically regulating the activity of an individual polymerase, phosphorylation at these sites might permanently assign a function and establish pools of specialized polymerases. Phosphorylation indirectly controls genome replication by regulating RNP assembly, and we now show that modifications on the viral polymerase directly control its actively to regulate product output.

## Materials and Methods

### Cells, viruses, plasmids, and transfections

All experiments were conducted with A549 (CCL-185), HEK 293T (CRL-3216), MDCK (CCCL-34), or MDBK (CCL-22) cells acquired from ATCC. Cells were maintained in Dulbeco’s modified Eagle’s medium (DMEM; Mediatech 10-013-CV) with 10% FBS and grown at 37°C in 5% CO_2_. Cells were regularly verified as mycoplasma negative using MycoAlert (Lonza LT07-218).

All virus and virus-derived protein expression constructs are based on A/WSN/1933. Expression constructs for the viral polymerase and NP were described previously (Engelhardt et al., 2005; Mehle and Doudna, 2008). FLAG-tagged PB1 expression constructs was generated via restriction cloning to express PB1 with a C-terminal 3X FLAG tag. Mutations were introduced by inverse PCR and confirmed by sequencing. The catalytically dead PB1a (PB1 D445A/D446A) and transcription-defective PB1 K669A/R670A mutants were previously characterized (Kerry et al., 2008; Vreede et al., 2004). Plasmid expressing the catalytic domain of PKCδ was previously described (Soh and Weinstein, 2003) (Addgene plasmid #16388).

Viruses were prepared using the pBD bi-directional reverse genetics system and pTM-All derivatives where multiple gene segments are consolidated on a single plasmid (Mehle and Doudna, 2008; Neumann et al., 2005). Rescued viruses were amplified on MDBK cells and titered on MDCK cells by plaque assay. When preparing mutant virus, the presence of the intended mutation was confirmed by sequencing RT-PCR product. WSN virus encoding FLAG-tagged PB2 (Dos Santos Afonso et al., 2005) was used for infections for mass spectrometric analysis and cRNA stabilization assays.

Transfections were performed using either TransIT 2020 (Mirus MIR5400) or PEI MAX40 (Polysciences 24765-1) following the manufacturers’ recommendation.

### Antibodies

FLAG-PB2 purifications for mass spectrometry were performed using M2 antibody (Sigma F1804) and captured using protein A dynabeads (Invitrogen 10002). Other FLAG immunoprecipitatons were performed with M2 Affinity Gel (Sigma A2220). The following antibodies were used for western blot analysis: α-PB1 (Mehle and Doudna, 2008), α-PB2 (Mehle and Doudna, 2008), α-PA (Genetex 125933), HA-HRP (3F10; Sigma 12013819001), M2-HRP (Sigma 8592). The mouse α-NP monoclonal antibody H16-L10-4R5 (Bioxcell BE0159) (Yewdell et al., 1981) was used for both immunoprecipitation and western blot analysis of NP.

### Synchronized single-cycle infections and multicycle replication assays

Viral infections with A549 cells were performed in virus growth medium (DMEM, 0.2% BSA, 25mM HEPES, 0.25 μg/mL TPCK-trypsin). Viral infections with 293T cells were performed in OptiMEM (Invitrogen) containing 2% FBS. For synchronized infections, cell monolayers were washed twice with ice-chilled PBS, incubated with inoculum for 1 hr at 4° C, followed by removal of inoculum and addition of pre-warmed fresh VGM (37° C) (Larson et al., 2019). Cells were maintained at 37° C for the duration of the infection. Multicycle replication assays were performed in A549 cells by inoculating cells in triplicate at an MOI 0.001. Virus was sampled at the indicated time points and titers were determined by plaque assay on MDCK cells.

Gene expression was measured during an asynchronous infection by inoculating A549 cells at an MOI of 0.05 for 8 h. Infections were terminated and total RNA was extracted from cells using TRIzol (Invitrogen). 250 ng of total RNA was subject to poly-dT primed reverse transcription using MMLV-RT. Resulting cDNA was used for qPCR to detect GAPDH and NP mRNA with the iTaq Universal SYBR Green Supermix (Bio-Rad 1725121). Fold changes in NP mRNA were calculated using the ΔΔC_T_ method.

### Polymerase activity assays

HEK 293T cells were transfected with plasmids expressing NP, PB2, FLAG-tagged PB1, FLAG-tagged PA, and segment 6 vRNA (NA). Total RNA was extracted using TRIzol 24 hr after transfection.

### Primer Extension analysis

RNA was subject to primer extension analysis for genomic (vRNA), complementary (cRNA), and messenger (mRNA) corresponding to segment 6. Primer extension analysis was performed as previously described for full-length and short (NP77) templates (Baker et al., 2018; Kirui et al., 2016). Briefly, RNA and the appropriate radiolabeled primers were boiled for 2 min and snap-chilled on ice. Samples were pre-heated to 42° C and pre-heated reaction mixture was added for a final reaction containing 50 mM Tris-HCl (pH 8.3), 75 mM KCl, 3 mM MgCl_2_, 5 mM DTT, 40 units RNAsin+ (Promega N2611), and house-made MMLV-RT (Kirui et al., 2016). Samples were incubated for 1 hr at 42° C. Reactions were terminated with an equal volume of 2x RNA loading dye (47.5% formamide, 0.01% SDS, 0.5 mM EDTA containing bromophenol blue and xylene cyanol), boiled for 2 mins and snap-chilled on ice prior to resolving on 6% (full-length templates) or 12% (short templates) denaturing polyacrylamide gels containing 0.5X TBE and 7M Urea. Gels were fixed (40% methanol, 10% acetic acid, 5% glycerol) for 30 mins, dried, quantified by phosphorimaging, and analyzed using Image Studio software (Licor).

### Preparation of samples for mass spectrometry

Mass spectrometry was performed on both transfected and infected cells. HEK 293T cells (3 × 100mm dishes containing approximately 6×10^6^ cells) were transfected to express PB1, FLAG-tagged PB2, PA and vNP76 (Turrell et al., 2013). 24 hr later, cells were lysed in RIPA Buffer (150mM NaCl, 50mM Tris pH 7.5, 0.5% w/v sodium deoxycholate, 0.1% w/v SDS, 1% w/v Ipegal 630, 2mM EDTA) in the presence of protease inhibitors and phosphatase inhibitors for 20 mins at 4°C. Lysates were sonicated and then cleared via centrifugation at 4° C. FLAG-tagged PB2 was immunoprecipated using M2 antibody overnight at 4° C. Immunocomplexes were captured using protein A dynabeads (Invitrogen 10002D) for 2 hr. Immunoprecipitations were washed twice with RIPA buffer and 4 times with NTE (100mM NaCl, 10mM Tris pH 7.5, 1mM EDTA). Protein was eluted with 8M urea in NTE. Samples were frozen at −80° C prior to mass spectrometric analysis.

For samples prepared from virally-infected cells, A549 cells were synchronously infected with WSN at an MOI of 5. Cells were collected at 2.5 hpi (30×10^6^ cells), 5 hpi (18×10^6^ cells), 7.5 hpi (18×10^6^ cells), and 10 hpi (12×10^6^ cells). Cell numbers were adjusted in an attempt to account for lower amounts of viral proteins early during infection. Cells were scraped into ice-chilled PBS and collected by centrifugation. NP immunoprecipitations were performed as above using α-NP monoclonal antibody H16-L10-4R5. Samples were also prepared using the same approach in separate experiments where A549 cells were infected with WSN at an MOI of 0.1 for 24 hr or an MOI of 5 for 6hr. Cell pellets and immunoprecipitations were frozen at −80° C prior to mass spectrometric analysis.

### Mass spectrometry

#### Sample preparation for Nano-LC-MS/MS

Infected cells were lysed in 6M guanidine-HCl for 10 min at 100 °C. Protein was precipitated by addition of methanol to a final concentration of 90% and pelleted by centrifugation at 12,000 x G for 10 min. The supernatant was discarded and pellet was resuspended in 8M urea, 50 mM Tris (pH 8.0), 10 mM tris(2-carboxyethyl)phosphine) (TCEP) and 40 mM chloroacetamide and rocked at room temperature for 30 min to reduce and alkylate cysteines. The sample was diluted to a urea concentration of less than 1.5 M with 50 mM Tris (pH 8.0) before adding protease grade trypsin (Promega) at an enzyme:protein ratio of 1:50 (mg:mg). The samples were rocked overnight at room temperature during the digestion. 10% trifluoroacetic acid was added to the solution to bring the pH of the sample less than 2 before desalting and peptide isolation using Strata-X reverse phase resin (Phenomenex). Sample were dried under reduced pressure, resuspended in 0.2% formic acid, and quantified by Pierce Quantiative Colorimetric Peptide Assay (Thermo Fisher Scientific). Phosphopeptides were enriched for each sample from 2 mg tryptic peptides using immobilized metal affinity chromatography (Ti-IMAC MagResyn, ReSyn Biosciences).

#### Nano-LC-MS/MS Data Acquisition

Each sample was analyzed using an Q-LTQ-OT tribrid mass spectrometer (Orbitrap Fusion Lumos) during a 90 min nano-liquid chromatography using a Dionex UltiMate 3000 RSLCnano system (Thermo Fisher Scientific). MS parameters differed for the analysis of phosphopeptides enriched and unenriched sample. For unenriched sample, MS1 survey scans were acquired in the Orbitrap (Resolution – 240K, AGC Target – 1×10^6^, Scan Range – 300-1,350 Da, Maximum Injection Time – 100 ms). MS2 spectra of observed precursors were acquired in the ion trap (Resolution – Rapid, AGC target 4×10^4^, Scan Range – 200-1,200, Maximum Injection Time – 18 ms) following quadrupole isolation (0.7 Da) and higher energy collisional dissociation (25% NCE). For phosphopeptides enriched samples, MS1 survey scans were acquired in the Orbitrap (Resolution – 60K, AGC Target – 1×10^6^, Scan Range – 300-1,350 Da, Maximum Injection Time – 50 ms). Observed precursors were also analyzed in the Orbitrap (Resolution – 15K, AGC Target – 5×10^4^, Scan Range – 150-1,500, Maximum Injection Time – 50 ms) following quadrupole isolation (1.6 Da) and higher energy collisional dissociation (25 % NCE). Monoisotopic precursor isolation and a dynamic exclusion of 15 s were enabled for both methods.

#### Data Analysis

Thermo RAW data files were searched using MaxQuant (version 1.5.3.51) with the Andromeda search algorithm against a concatenated target-decoy database of human and influenza proteins using default search tolerances (Cox et al., 2011; Elias and Gygi, 2007). Specified search parameters included the fixed modification of carbamidomethylation at cysteine residues and variable modification for methinine oxidation. Phosphorylation of serine, threonine, and tyrosine were specified as variable modifications for phosphopeptide enriched data. Label free quantitation and intensity based absolute quantitation were enabled (Cox et al., 2014; Schwanhäusser et al., 2011).

### Polymerase formation assays

Polymerase assembly was measured as before (Kirui et al., 2014). FLAG-tagged PB1, PB2, and PA were expressed in transfected HEK 293T cells for 48 hr. Cells were lysed in co-IP buffer (50mM Tris pH 7.4, 150mM NaCl, 0.5% Igepal CA-630) in the presence of protease inhibitors for 20mins at 4° C. Lysates were clarified by centrifugation and pre-cleared with protein-A agarose (Santa Cruz Biotech sc-2001) for 1hr. Lysates were then transferred to a new microcentrifuge tube and BSA was added to a final concentration of 5 mg/ml. FLAG-PB1 was immunoprecipitated overnight with M2-agarose. Immunoprecipitations were washed twice with co-IP buffer containing 5 mg/mL BSA and 500mM NaCl and twice with co-IP buffer. Bound proteins were eluted by boiling in Laemmli buffer. Samples were then assayed via western blot analysis for presence of PB1, PB2, and PA.

### RNP reconstitution assays

NP, PB2, FLAG-tagged PB1, FLAG-tagged PA, and segment 6 vRNA (NA) were expressed in transfected HEK 293T cells for 48 hr, following prior approaches (Baker et al., 2018). Cells were lysed in co-IP buffer in the presence of protease inhibitors. Lysates were clarified by centrifugation, pre-cleared protein A agarose (Santa Cruz Biotech sc-2001) for 1 hr, and transferred to a new tube where BSA was added to a final concentration of 5 mg/mL. NP was immunoprecipitated overnight with 3 µg anti-NP antibody. Immunocomplexes were captured using protein A agarose (Sigma P2545) for 1 hr, washed twice with co-IP buffer containing 5 mg/mL BSA and 500 mM NaCl, and twice with co-IP buffer. Bound proteins were eluted by boiling in Laemmli buffer. Samples were then assayed via western blot analysis for presence of NP, PB1, and PA.

### cRNA stabilization assay

cRNA stabilization was measured as previously described (Vreede et al., 2004, 2011). Briefly, HEK 293T cells were transfected to express the viral polymerase with the indicated PB1 subunit and an oligomerization deficient NP (NP_E339A_). 24 hr post-transfection, cells were treated with actinomycin D (5 µg/mL) (Sigma A1410) for 30 mins prior to asynchronous infection with WSN in the presence of actinomycin D. Cells were harvested 6 hpi. Total RNA was extracted using TRIzol and used in primer extension analysis.

### RNA immunoprecipitation vRNA binding assay

RNPs with FLAG-tagged PB1 were reconstituted in HEK 293T cells as above. Cells were lysed 48 hr post-transfection in co-IP buffer supplemented with both protease inhibitors and RNAsin (Promega N2515, 100 units/mL). Lysates were processed and immunoprecipitations were performed as described above for the polymerase formation assay. Protein from 10% of the immunoprecipitate was eluted by boiling in Laemmli buffer and assayed via western blot. RNA from 90% of the immunoprecipitate was extracted using TRIzol and analyzed by primer extension.

### Statistics

Data represent at least 2-3 independent biological replicates. Technical replicates are indicated for each figure. Quantitative data are shown as mean ± standard deviation for one biological replicate or the mean of means ± standard error of measurement for multiple biological replicates. Single pair-wise comparisons were analyzed by Student’s t-test. Multiple comparisons were performed by a one-way ANOVA followed by Dunnett’s *post hoc* analysis of pair-wise comparisons to WT. P<0.05 was considered significant. Statistic were calculated in Prism 8.

## Acknowledgements

We thank members of the Mehle and Coon lab for their constructive input. This work is funded by R01AI125271 to AM and JJC and R35GM118110 to JJC. ARD is funded by T32AI078985. GMW is supported by T32GM008349. AM holds an Investigators in the Pathogenesis of Infectious Disease Award from the Burroughs Wellcome Fund.

---

**Figure 4 Figure Supplement 1:**
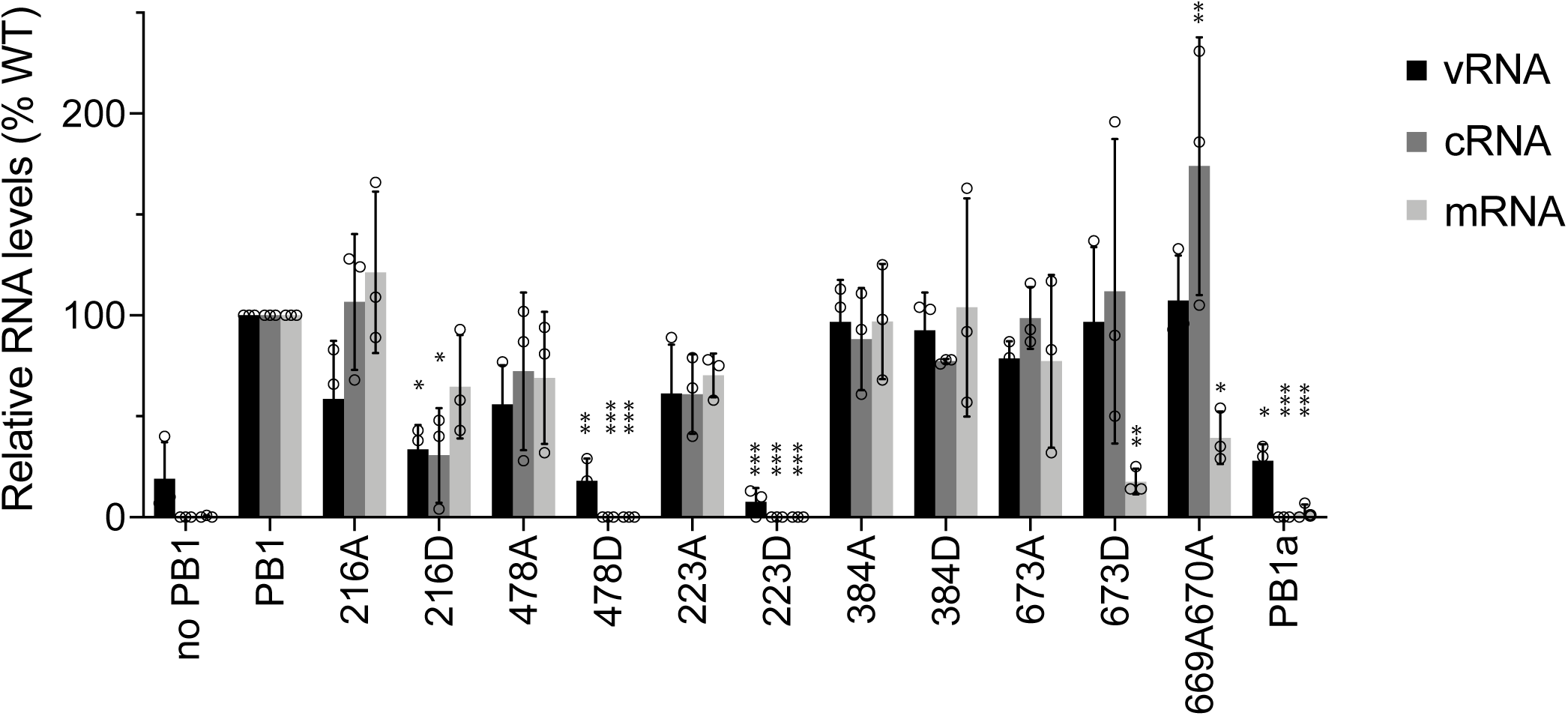
Quantification of replicate primer extension assays. Three independent primer extension assays were quantified with each RNA species normalized to WT within an experiment. Data are presented at mean ± sd. * < 0.05, ** < 0.01, *** < 0.001 = P for one-way ANOVA with Dunnett’s post hoc compared to WT.

**Figure 5 Figure Supplement 1:**
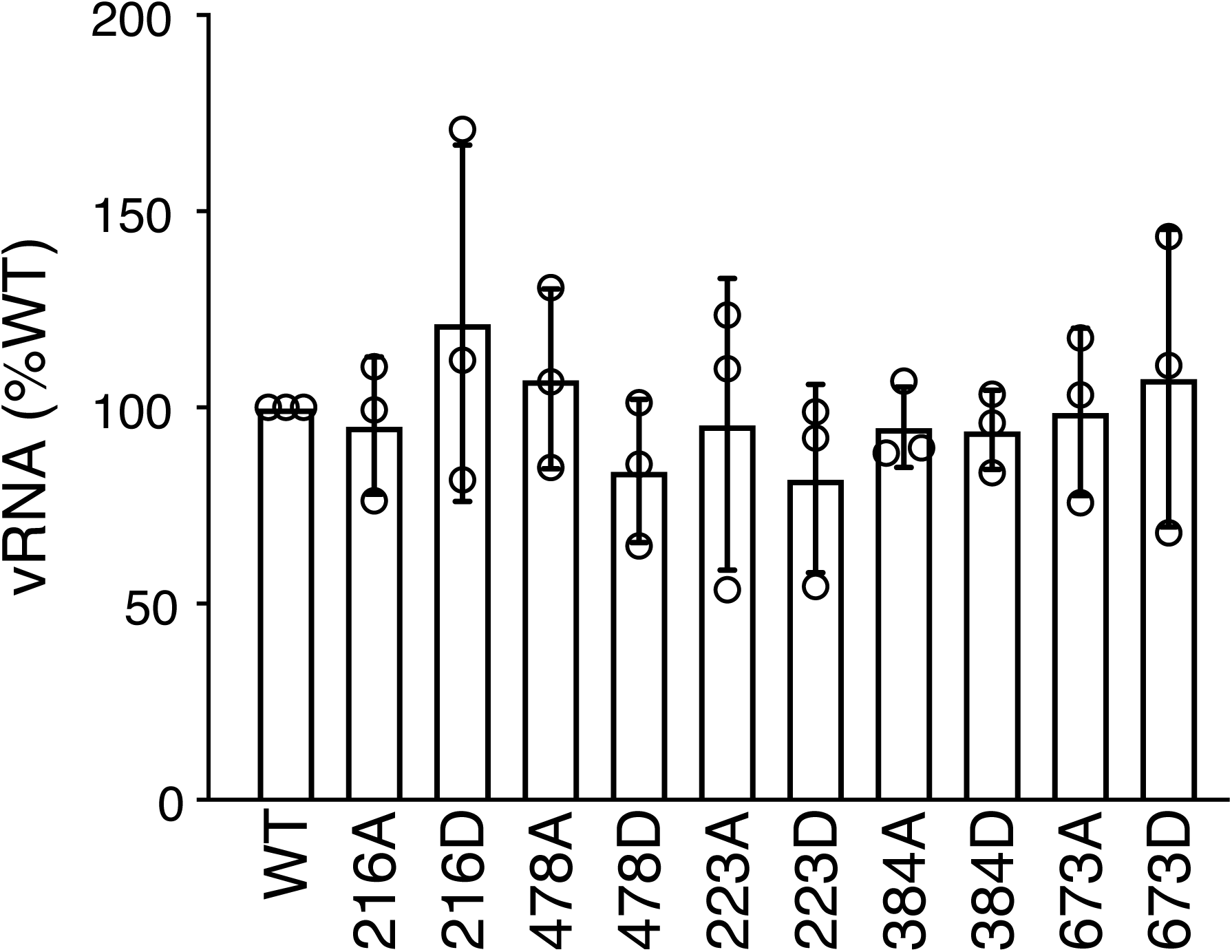
Quantification of replicate cRNA stabilization assays. vRNA levels were quantified from three independent stabilization assays and normalized to WT. Data are presented at mean ± sd. There was no significant difference when analyzed by a one-way ANOVA.

## References

Arranz, R., Coloma, R., Chichon, F.J., Conesa, J.J., Carrascosa, J.L., Valpuesta, J.M., Ortin, J., and Martin-Benito, J. (2012). The structure of native influenza virion ribonucleoproteins. Science (80-.). 338, 1634–1637.

Baker, S.F., Ledwith, M.P., and Mehle, A. (2018). Differential Splicing of ANP32A in Birds Alters Its Ability to Stimulate RNA Synthesis by Restricted Influenza Polymerase. Cell Rep. 24, 2581-2588.e4.

Biedenkopf, N., Lier, C., and Becker, S. (2016). Dynamic Phosphorylation of VP30 Is Essential for Ebola Virus Life Cycle. J. Virol. 90, 4914–4925.

Bouloy, M., Plotch, S.J., and Krug, R.M. (1978). Globin mRNAs are primers for the transcription of influenza viral RNA in vitro. Proc. Natl. Acad. Sci. 75, 4886–4890.

Cartee, T.L., and Wertz, G.W. (2001). Respiratory Syncytial Virus M2-1 Protein Requires Phosphorylation for Efficient Function and Binds Viral RNA during Infection. J. Virol. 75, 12188 LP – 12197.

Chen, L., Zhang, S., Banerjee, A.K., and Chen, M. (2013). N-terminal phosphorylation of phosphoprotein of vesicular stomatitis virus is required for preventing nucleoprotein from binding to cellular RNAs and for functional template formation. J. Virol. 87, 3177–3186.

Cox, J., Neuhauser, N., Michalski, A., Scheltema, R.A., Olsen, J. V., and Mann, M. (2011). Andromeda: A peptide search engine integrated into the MaxQuant environment. J. Proteome Res. 10, 1794–1805.

Cox, J., Hein, M.Y., Luber, C.A., Paron, I., Nagaraj, N., and Mann, M. (2014). Accurate proteome-wide label-free quantification by delayed normalization and maximal peptide ratio extraction, termed MaxLFQ. Mol. Cell. Proteomics 13, 2513–2526.

Cui, L., Mahesutihan, M., Zheng, W., Meng, L., Fan, W., Li, J., Ye, X., Liu, W., and Sun, L. (2018). CDC25B promotes influenza A virus replication by regulating the phosphorylation of nucleoprotein. Virology 525, 40–47.

Dawson, A.R., and Mehle, A. (2018). Flu’s cues: Exploiting host post-translational modifications to direct the influenza virus replication cycle. PLoS Pathog. 14, e1007205.

Deng, T., Sharps, J., Fodor, E., and Brownlee, G.G. (2005). In vitro assembly of PB2 with a PB1-PA dimer supports a new model of assembly of influenza A virus polymerase subunits into a functional trimeric complex. J. Virol. 79, 8669–8674.

Elias, J.E., and Gygi, S.P. (2007). Target-decoy search strategy for increased confidence in large-scale protein identifications by mass spectrometry. Nat. Methods 4, 207–214.

Engelhardt, O.G., Smith, M., and Fodor, E. (2005). Association of the influenza A virus RNA-dependent RNA polymerase with cellular RNA polymerase II. J. Virol. 79, 5812–5818.

Fodor, E., and te Velthuis, A.J.W. (2019). Structure and Function of the Influenza Virus Transcription and Replication Machinery. Cold Spring Harb. Perspect. Med.

Hay, A.J., Lomniczi, B., Bellamy, A.R., and Skehel, J.J. (1977). Transcription of the influenza virus genome. Virology 83, 337–355.

Herz, C., Stavnezer, E., Krug, R.M., and Gurney, T. (1981). Influenza virus, an RNA virus, synthesizes its messenger RNA in the nucleus of infected cells. Cell 26, 391–400.

Hutchinson, E.C., Denham, E.M., Thomas, B., Trudgian, D.C., Hester, S.S., Ridlova, G., York, A., Turrell, L., and Fodor, E. (2012). Mapping the Phosphoproteome of Influenza A and B Viruses by Mass Spectrometry. PLoS Pathog. 8.

Jackson, D.A., Caton, A.J., McCready, S.J., and Cook, P.R. (1982). Influenza virus RNA is synthesized at fixed sites in the nucleus. Nature 296, 366–368.

Kerry, P.S., Willsher, N., and Fodor, E. (2008). A cluster of conserved basic amino acids near the C-terminus of the PB1 subunit of the influenza virus RNA polymerase is involved in the regulation of viral transcription. Virology 373, 202–210.

Kirui, J., Bucci, M.D., Poole, D.S., and Mehle, A. (2014). Conserved features of the PB2 627 domain impact influenza virus polymerase function and replication. J. Virol. 88, 5977–5986.

Kirui, J., Mondal, A., and Mehle, A. (2016). Ubiquitination up-regulates influenza virus polymerase function. J. Virol. 90, 10906–10914.

Klumpp, K., Ruigrok, R.W., and Baudin, F. (1997). Roles of the influenza virus polymerase and nucleoprotein in forming a functional RNP structure. EMBO J 16, 1248–1257.

Kouba, T., Drncová, P., and Cusack, S. (2019). Structural snapshots of actively transcribing influenza polymerase. Nat. Struct. Mol. Biol. 26, 460.

Larson, G.P., Tran, V., Yú, S., Caì, Y., Higgins, C.A., Smith, D.M., Baker, S.F., Radoshitzky, S.R., Kuhn, J.H., and Mehle, A. (2019). EPS8 Facilitates Uncoating of Influenza A Virus. Cell Rep. 29, 2175–2183.

Mehle, A., and Doudna, J.A. (2008). Inhibitory activity restricts the function of an avian-like influenza polymerase in primate cells. Cell Host Microbe 4, 111–122.

Moeller, A., Kirchdoerfer, R.N., Potter, C.S., Carragher, B., and Wilson, I.A. (2012). Organization of the influenza virus replication machinery. Science (80-.). 338, 1631–1634.

Mondal, A., Victor, K.G., Pudupakam, R.S., Lyons, C.E., and Wertz, G.W. (2014). Newly Identified Phosphorylation Site in the Vesicular Stomatitis Virus P Protein Is Required for Viral RNA Synthesis. J. Virol. 88, 1461–1472.

Mondal, A., Potts, G.K., Dawson, A.R., Coon, J.J., and Mehle, A. (2015). Phosphorylation at the homotypic interface regulates nucleoprotein oligomerization and assembly of the influenza virus replication machinery. PLoS Pathog. 11, e1004826.

Mondal, A., Dawson, A.R., Potts, G.K., Freiberger, E.C., Baker, S.F., Moser, L.A., Bernard, K.A., Coon, J.J., and Mehle, A. (2017). Influenza virus recruits host protein kinase C to control assembly and activity of its replication machinery. Elife 6, e26910.

Neumann, G., Fujii, K., Kino, Y., and Kawaoka, Y. (2005). An improved reverse genetics system for influenza A virus generation and its implications for vaccine production. Proc. Natl. Acad. Sci. U. S. A. 102, 16825–16829.

Perez, J.T., Varble, A., Sachidanandam, R., Zlatev, I., Manoharan, M., García-Sastre, A., and TenOever, B.R. (2010). Influenza A virus-generated small RNAs regulate the switch from transcription to replication. Proc. Natl. Acad. Sci. U. S. A.

Pflug, A., Guilligay, D., Reich, S., and Cusack, S. (2014). Structure of influenza A polymerase bound to the viral RNA promoter. Nature 516, 355–360.

Plotch, S.J., Bouloy, M., and Krug, R.M. (1979). Transfer of 5’-terminal cap of globin mRNA to influenza viral complementary RNA during transcription in vitro. Proc. Natl. Acad. Sci. 76, 1618–1622.

Pons, M.W., Schulze, I.T., Hirst, G.K., and Hauser, R. (1969). Isolation and characterization of the ribonucleoprotein of influenza virus. Virology 39, 250–259.

Reich, S., Guilligay, D., Pflug, A., Malet, H., Berger, I., Crepin, T., Hart, D., Lunardi, T., Nanao, M., Ruigrok, R.W., et al. (2014). Structural insight into cap-snatching and RNA synthesis by influenza polymerase. Nature 516, 361–366.

Reich, S., Guilligay, D., and Cusack, S. (2017). An in vitro fluorescence based study of initiation of RNA synthesis by influenza B polymerase. Nucleic Acids Res. 45, 3353–3368.

Richard, C.-A., Rincheval, V., Lassoued, S., Fix, J., Cardone, C., Esneau, C., Nekhai, S., Galloux, M., Rameix-Welti, M.-A., Sizun, C., et al. (2018). RSV hijacks cellular protein phosphatase 1 to regulate M2-1 phosphorylation and viral transcription. PLOS Pathog. 14, e1006920.

Robb, N.C., Smith, M., Vreede, F.T., and Fodor, E. (2009). NS2/NEP protein regulates transcription and replication of the influenza virus RNA genome. J. Gen. Virol. 90, 1398–1407.

Santos Afonso, E., Escriou, N., Leclercq, I., van der Werf, S., and Naffakh, N. (2005). The generation of recombinant influenza A viruses expressing a PB2 fusion protein requires the conservation of a packaging signal overlapping the coding and noncoding regions at the 5′ end of the PB2 segment. Virology 341, 34–46.

Schwanhäusser, B., Busse, D., Li, N., Dittmar, G., Schuchhardt, J., Wolf, J., Chen, W., and Selbach, M. (2011). Global quantification of mammalian gene expression control. Nature 473, 337–342.

Soh, J.-W., and Weinstein, I.B. (2003). Roles of Specific Isoforms of Protein Kinase C in the Transcriptional Control of Cyclin D1 and Related Genes. J. Biol. Chem. 278, 34709–34716.

Tigabu, B., Ramanathan, P., Ivanov, A., Lin, X., Ilinykh, P.A., Parry, C.S., Freiberg, A.N., Nekhai, S., and Bukreyev, A. (2018). Phosphorylated VP30 of Marburg Virus Is a Repressor of Transcription. J. Virol. 92, e00426–18.

Turrell, L., Lyall, J.W., Tiley, L.S., Fodor, E., and Vreede, F.T. (2013). The role and assembly mechanism of nucleoprotein in influenza A virus ribonucleoprotein complexes. Nat. Commun. 4, 1511–1591.

Vreede, F.T., Jung, T.E., and Brownlee, G.G. (2004). Model Suggesting that Replication of Influenza Virus Is Regulated by Stabilization of Replicative Intermediates. J. Virol. 78, 9568–9572.

Vreede, F.T., Ng, A.K.-L., Shaw, P.-C., and Fodor, E. (2011). Stabilization of Influenza Virus Replication Intermediates Is Dependent on the RNA-Binding but Not the Homo-Oligomerization Activity of the Viral Nucleoprotein. J. Virol. 85, 12073–12078.

Weber, A., Dam, S., Saul, V. V., Kuznetsova, I., Müller, C., Fritz-Wolf, K., Becker, K., Linne, U., Gu, H., Stokes, M.P., et al. (2019). Phosphoproteome Analysis of Cells Infected with Adapted and Nonadapted Influenza A Virus Reveals Novel Pro- and Antiviral Signaling Networks. J. Virol. 93.

Yewdell, J.W., Frank, E., and Gerhard, W. (1981). Expression of influenza A virus internal antigens on the surface of infected P815 cells. J. Immunol. 126, 1814–1819.

York, A., Hengrung, N., Vreede, F.T., Huiskonen, J.T., and Fodor, E. (2013). Isolation and characterization of the positive-sense replicative intermediate of a negative-strand RNA virus. Proc. Natl. Acad. Sci. U. S. A. 110, E4238–45.

